# Model-based ordination of pin-point cover data: effect of management on dry heathland

**DOI:** 10.1101/2020.03.05.980060

**Authors:** Christian Damgaard, Rikke Reisner Hansen, Francis K. C. Hui

## Abstract

Recently, there has been an increasing interest in model-based approaches for the statistical modelling of the joint distribution of multi-species abundances. The Dirichlet-multinomial distribution has been proposed as a suitable candidate distribution for the joint species distribution of pin-point plant cover data and is here applied in a model-based ordination framework. Unlike most model-based ordination methods, both fixed and random effects are in our proposed model structured as *p*-dimensional vectors and added to the latent variables before the inner product with the species-specific coefficients. This changes the interpretation of the parameters, so that the fixed and random effects now measure the relative displacement of the vegetation by the fixed and random factors in the *p*-dimensional latent variable space. This parameterization allows statistical inference of the effect of fixed and random factors in vector space, and makes it easier for practitioners to perform inferences on species composition in a multivariate setting. The method was applied on plant pin-point cover data from dry heathlands that had received different management treatments (burned, grazed, harvested, unmanaged), and it was found that treatment have a significant effect on heathland vegetation both when considering plant functional groups or when the taxonomic resolution was at the species level.

## Introduction

The statistical treatment of whole communities with multiple species has traditionally relied on ordination methods that use distance measures to reduce the plot by species abundance matrix into a low dimensional vector of distances among plots. Traditional methods for ordination, such as Non Metric Multidimensional scaling (NMDS) or Principal Component Analysis (PCoA), rely on strictly algorithmic approaches with no underlying statistical model of species abundance (McCune and Grace 2002). To accommodate potential properties inherent in the data (e.g. a strong mean-variance relationship with count data), these methods may apply a variety of distance measures or data transformations. However, commonly applied distance metrics, such as Euclidean, Manhattan, or Bray–Curtis distance, often make implicit assumptions on the mean-variance relationship that may not be entirely compatible with the data at hand. This can result in potentially incorrect conclusions, e.g. location effects may be confounded within the dispersion effects (Warton et al. 2012, Warton and Hui 2017).

Recently, a number of ordination techniques, where the distribution of species abundance is explicitly taken into account, have been developed. Collectively, these new techniques are referred to as model-based ordination. In univariate statistics, for instance, these issues of mean-variance relations in the species response have long been addressed with generalized linear models (GLM’s) and their mixed model counterparts, where the mean-variance relationship is modelled for each response variable (Bolker et al. 2009). Model-based ordination can thus be regarded as multivariate extensions of GLM’s. This method offers the possibility of adjusting the distribution family to e.g. negative binomial distribution for overdispersed count data and the Bernoulli distribution for presence-absence data to better account for the inherent mean-variance relationship in the species observations (Hui et al. 2015, Warton et al. 2015). Like GLM’s, model-based ordination methods further allow for a check of the validity of such model assumptions. Through row/column standardizations, as incorporated in model-based approaches, each taxon or species contribution to the sample location in ordination space is accounted for (Hawinkel et al. 2019) as well as potential clustering of data across different levels of study structure. For instance, hierarchical structures caused by host-parasite dynamics and reflecting nested design experiments (Björk et al. 2018). Hence, additions to the model-based ordination literature are continually emerging (e.g. Sohn and Li 2018, Hawinkel et al. 2019, Niku et al. 2019).

Arguably, the most popular method of model-based ordination that has emerged is latent variable modelling, which involves modelling community composition through a small set of underlying latent variables that reduce the dimension of the species abundance data from the number of species to the dimension of the latent variables (Hui et al. 2015, Warton et al. 2015, Warton et al. 2016, Niku et al. 2017). In doing so, the latent variables also express the residual variation owing to factors beyond those of the measured predictors included in the model, e.g. possible biotic interactions or phylogenetic relatedness between species. More specifically, in latent variable modelling the species composition of each plot is described by a vector with dimension *p* that is much less than the number of species. These latent variables are then fitted to the species abundance data using the relevant distribution of species abundance. Since the latent variables are not observed, they are treated as random effects, meaning that they need to be predicted at the same time that coefficients associated with them are estimated (often called loadings) along with other potential site, treatment, and environmental effects.

Ecological multivariate data analyzed using these methods often presents as count data or presence-absence data. Because plants are sessile, they facilitate registration of multiple characteristics related to vegetation structure and microclimate. The most common way to quantify plant species abundance in light-open plant communities is to measure the cover, which is the relative area of the species when projected onto the surface. Unlike plant counts or density, plant cover takes the size of individuals into account. A relatively objective method for measuring plant cover is the pin-point (or point-intercept) method in a frame or along a line, which has been widely employed in the field. This involves vertically inserting a thin pin into the vegetation a number of times in a fixed design, and the cover of a species is measured by the proportion of the inserted pins that touches the species (Lindquist 1931, Levy and Madden 1933, Damgaard and Irvine 2019). The pin-point method is not relevant for measuring the abundance of rare species and has been shown to underestimate species richness (Bråkenhielm and Qinghong 1995). Importantly, the pin-point method allows unbiased aggregation of single species cover measures at the pin level into cover data for higher plant taxa, e.g. the cover of grasses or herbal plants.

Many plant species are spatially aggregated, and it is important to take this spatial aggregation into account when modelling multi-species pin-point plant cover data. Recently, a reparametrized Dirichlet-multinomial distribution has been proposed for modelling such cover data (Damgaard 2015, 2018). The Dirichlet distribution is the conjugate prior of the multinomial distribution and a multivariate generalization of the beta-binomial distribution, which has previously been used with success to model pin-point cover data of spatially aggregated plant species (Damgaard 2009, 2013, Damgaard and Irvine 2019). Moreover, the Dirichlet-multinomial distributed model has been applied for modelling multi-species vegetation dynamics in e.g. heathland ecosystems (Damgaard 2015, Damgaard et al. 2017, Damgaard 2019).

The aim of this study is two-fold; i) to modify the model-based ordination method previously suggested by among others Hui et al. (2015) so that it is applicable to multi-species pin-point cover data, and ii) to reparametrize the underlying latent variable model so that the effect of fixed and random factors on the species composition may be investigated and tested in the same underlying multidimensional space as the latent variables characterizing the ordination.

The proposed method will be illustrated on multi-species pin-point cover data from six Danish dry heathlands, where different nature management practices have been applied and, more specifically, it will be used to assess whether the form of management has had an effect on heathland vegetation.

## Model

The objective of the proposed model is to reduce a plant species abundance matrix with *k* plots and *n* species to a *k*\**p* matrix of latent variables, where *p* < *n* is the number of latent variables, and the plant abundance of each species is measured by its cover using the pin-point method. The effects of observed covariates, including e.g. the experimental unit and design effects on species composition, may be entered as fixed and/or random effects into the mean structure as well. The inclusion of such effects means that the latent variables then model the so-called residual correlation between species, i.e. any covariation that cannot be explained by the observed predictors and experimental design effects (Warton et al. 2015, Niku et al. 2017).

Partly following Hui et al. (2015) and adopting a Bayesian framework for estimation and inference, we propose that the mean cover of species *j* in plot *i*, denoted here as *q_ij_*, be modelled as follows

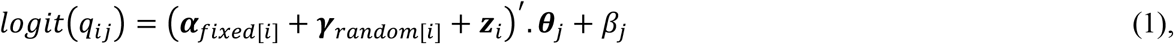

where all parameters in bold denote vectors of dimension *p*.

In this model, the vector ***α***_*fixed*[*i*]_ denotes a fixed effect applied to plot *i*, for instance a treatment effect, and are assigned weak prior distributions, e.g. *N_p_* (**0**,100 **I**), which is a p-dimensional multivariate normal distribution with zero mean vector and a covariance matrix with a diagonal matrix with all diagonal elements set to 100. Note that for reasons of parameter identifiability, it is assumed that these fixed effects are not unique to each plot (which is almost always the case).

Next, the vector ***γ***_*random*[*i*]_ model denotes a random effect applied to plot *i*, e.g. the location in a hierarchical experimental design, and are assumed to be drawn from a common multivariate normal distribution with a zero mean vector and an unstructured *p*\**p* covariance matrix, *N_p_*(**0, Σ**). For example, when *p* = 2, as is commonly considered for the purposes of ordination, 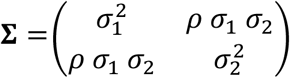. In this case, the standard deviations *σ_i_* are assigned a uniform positive prior, while the correlation coefficient *ρ* is assigned a uniform prior between −1 and 1. As with the fixed effects, the random effects are assumed not to be at plot level for reasons of parameter identifiability, e.g. in our case study the random effect is at the level of the site, in which multiple plots are nested. In the above notation (eq. 1), the model is fitted with only one fixed factor and one random factor, however, the model may be extended to include either more fixed or random effects, as well as including a spatial correlated random component. Conversely, the model may be simplified by either omitting the fixed and random effect.

Finally, the vector ***z**_i_* denotes a vector of *p* latent variables for plot *i* and, as is standard, is assumed to come from a standard normal distribution *N_p_* (**0,I**), where the zero mean vector and identity covariance matrix are used to avoid location and scale invariance and ensure that the parameters in the model are identifiable (Hui et al. 2015, Niku et al. 2017).

The vector *θ_j_* denotes a p-dimensional vector of coefficients or loading for species *j* (e.g. Hui et al. 2015) and is, similarly to the fixed effects above, assigned a weak prior *N_p_* (**0**,100 **I**). Note that in order to ensure that the parameters are identifiable, we further apply a standard constraint of assuming that the upper triangular portion of the *n* * *p* matrix of species-specific coefficients is constrained to be zero, while the diagonal elements are constrained to be positive (see Hui et al. 2015, Niku et al. 2017). The loadings can be interpreted as quantifying the species responses to the unmeasured latent variables and can also be plotted in conjunction with the ordinations as a modelbased biplot (e.g. Warton et al. 2015). Finally, the quantities *β_j_* denote a vector of constants chosen *a priori*, which are required to ensure that *logit*(*q_ij_*) is a real number.

The main difference between the present method and the original latent variable ordination approach proposed by, among others, Hui et al. (2015) and Warton et al. (2015) is that both the fixed and random effects, ***α***_*fixed*[*i*]_ and ***γ***_*random*[*i*]_ are vectors of dimension*p* and are added to the latent variables prior to their inner product with the species-specific coefficients. This changes the interpretation of the parameters so that the fixed and random effects now measure the relative displacement of the vegetation by the fixed and random factors in the *p*-dimensional latent space. Consequently, the latent variable ***z**_i_* may now more properly be referred to as modelling the residual correlation between species, i.e. the covariation after the fixed and random effects are taken into account. The scale of the displacement by the fixed and random effects is measured relatively to the residual latent variables, which are fixed to a unit standard variation for ease of interpretation (and parameter identifiability). This parameterization of the underlying latent variable model uses the concepts of fixed and random effects in an analogous way as to how the terms are used in generalized linear mixed effects models (Bolker et al. 2009). It is anticipated that this analogy will make it easier for practitioners to test e.g. treatment effects on species composition in a multivariate setting.

The underlying motivation for using model-based ordination instead of traditional ordination is to take the distributional properties of the species abundance sampling into account such that e.g. the correct mean-variance relationship is used in the modelling (e.g. Warton et al. 2012, Warton and Hui 2017). In this study, we consider the multi-species pin-point cover data, and it has previously been demonstrated that a reparametrized Dirichlet-multinomial distribution is a suitable candidate distribution to model multi-species pin-point cover data (Damgaard 2015, Damgaard et al. 2017, Damgaard 2019). The advantage of using the reparametrized Dirichlet-multinomial distribution is that the degree of spatial aggregation in plant communities is taken into account and explicitly modelled by a parameter *δ* (Damgaard 2018). More specifically, the observed hierarchical multi-species pin-point cover data, *Y*, is modelled by a mean cover vector of the *n*-1 first species, *q_j_*, and a parameter *δ*, which measures the degree of intra-specific spatial aggregation, by,

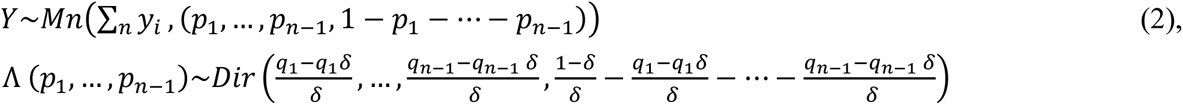

At the limit when *δ* → 0, the reparametrized Dirichlet-multinomial distribution degenerates into the multinomial distribution. Here, the prior distributions of *q_j_* and *δ* are assumed to be uniformly distributed between (0, 1) and (0.01, 0.95), respectively. For more details on the properties of the reparametrized Dirichlet-multinomial distribution, see Damgaard (2018).

### Estimation

The proposed latent variable model was estimated using Bayesian Markov Chain Monte Carlo (MCMC) methods using the Metropolis-Hastings algorithm with normally distributed candidate distributions. Specifically, we considered a MCMC chain with a burn-in period of 70,000 iterations followed by 30,000 additional iterations.

Trace plots of the sampling chains of all parameters and latent variables were examined to assess their mixing properties and convergence of the MCMC chain. Additionally, the overall fitting properties of the model were checked by examining the regularity and shape of the marginal distribution of parameters as well as the distribution of the deviance (= −2*log L*(*Y\θ*)). The efficiency of the MCMC procedure was assessed by inspecting the evolution in the deviance.

For ordination and when *p* = 2, we constructed plots of the posterior mean values of (***α***_*fixed*[*i*]_ + ***γ***_*random*[*i*]_ + ***z**_i_*) for each plot along with 95% credibility regions of ***α***_*fixed*[*i*]_, noting that these are p-dimensional regions rather than the unusual one-dimensional credibility intervals seen in mixed models and the standard latent variable ordination approach. These credibility regions enabled us to make statistical inferences on the effect of the fixed factors on species composition, e.g. if the 95% credibility regions of two fixed factors did not overlap. In such cases, we concluded that there is clear statistical evidence that the effects of the two factors on the species composition differ substantially. Consequently, the suggested parameterization in equation (1) will allow us to utilize the model-based ordination framework for testing purposes on the latent space. Furthermore, posterior means of the species coefficients, ***θ**_j_* for each cover class were also extracted and superimposed on top of the ordination diagram to visualize the contribution of each cover class (species) to the location of plots in the diagram, although note that the mean of the last species (*q_n_*) is not estimated in eqn. 2.

All calculations were done using *Mathematica* (Wolfram 2016).

### Case study

To assess the effect of heathland management on plant communities, plant species were registered using the pin-point method. During the fall of 2018, the cover of heathland plant species was registered at 6 heathland sites encompassing four different heathland management regimes: harvesting, grazing, burning and unmanaged, i.e. abandonment of management. At each site several of the four management regimes were applied. Within each site and management combination, the pin-point measurements were repeated in each of ten randomly positioned plots (Electronic Supplement, Appendix A). The pin-point frames had 25 pins within an area of 0.5m x 0.5m. Pinpoint data were then aggregated at the pin-level into five cover classes: Dwarf shrubs, graminoids, forbs, mosses and lichens. That is, if e.g. two graminoid species where hit by the same pin, they were aggregated to a single hit (Fig. 1).

**Fig. 1.**
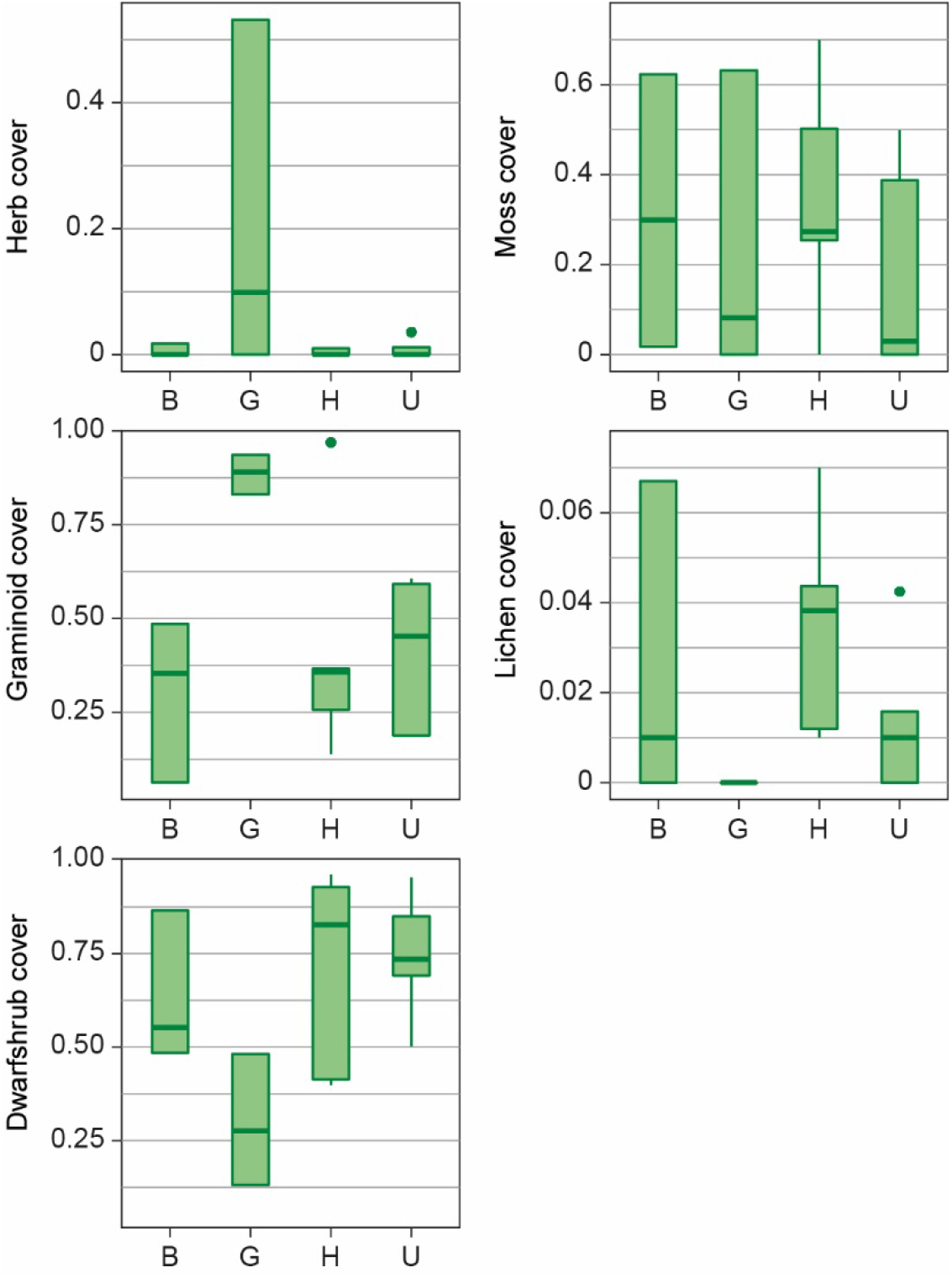
The cover of the aggregated species at the different management treatments B: burned, G: grazed, H: harvested, U: unmanaged.

We modelled both the aggregated pin-point data and the underlying single species pin-point data (number of species: 34) using the proposed latent variable ordination approach formulated above, where the four different heathland management regimes were assumed to be *p*-dimensional fixed effects, i.e. ***α**_treatment[i]_* in equation (1), and the site was assumed to be a *p*-dimensional random effect, i.e. ***γ***_*site*[*i*]_ in equation (1). For the purpose of visualization, the dimension of the latent variable model (*p*) was set to 2.

## Results

The MCMC iterations demonstrated fair mixing properties (Electronic supplement, Appendix B), and the marginal posterior distribution of selected parameters and compound parameters are summarized in Table 1 and Fig. 2.

**Table 1.**
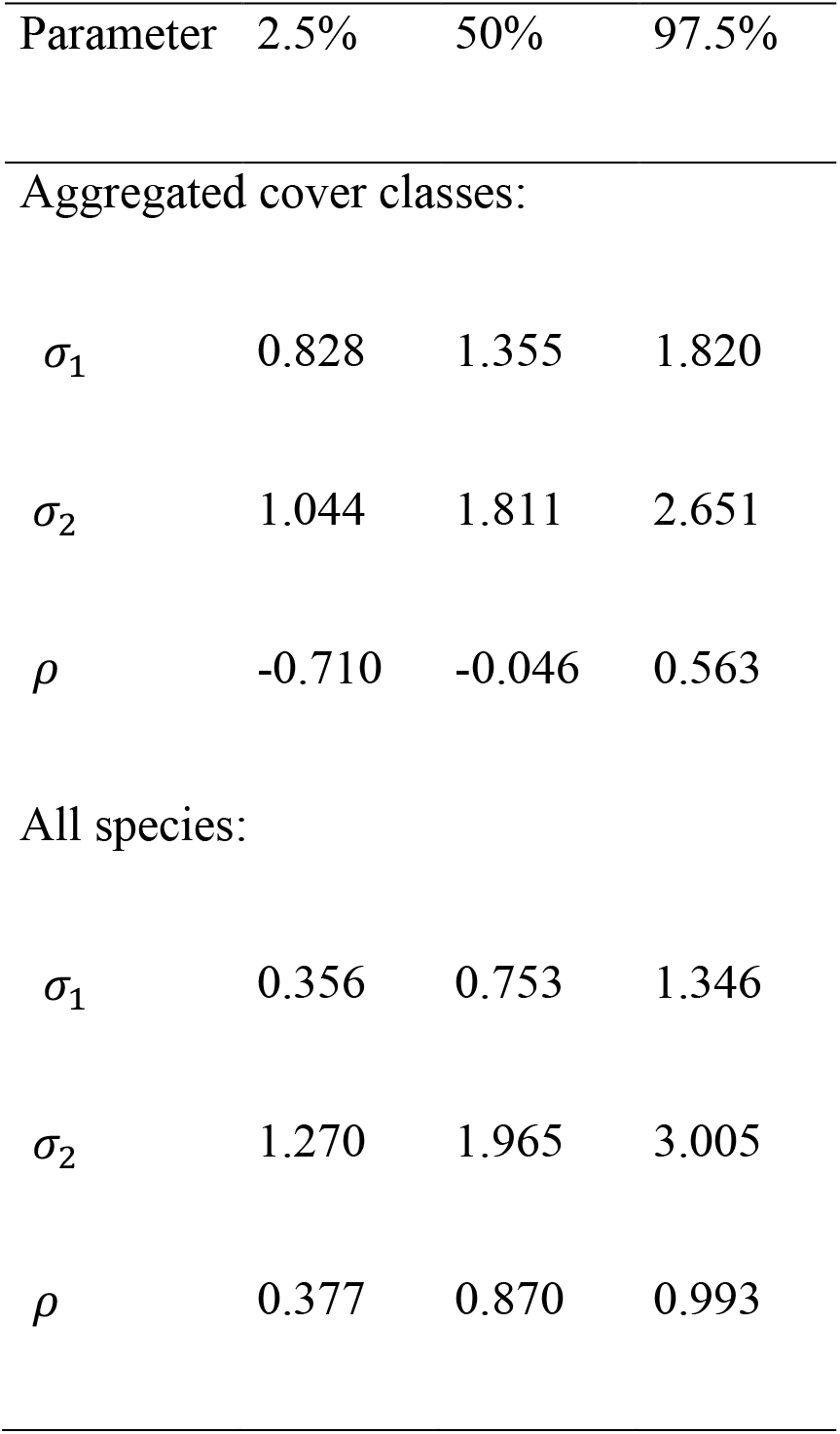
Marginal credible intervals for the posterior distribution of the parameters in 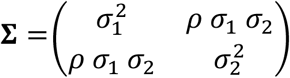

**Fig. 2.**
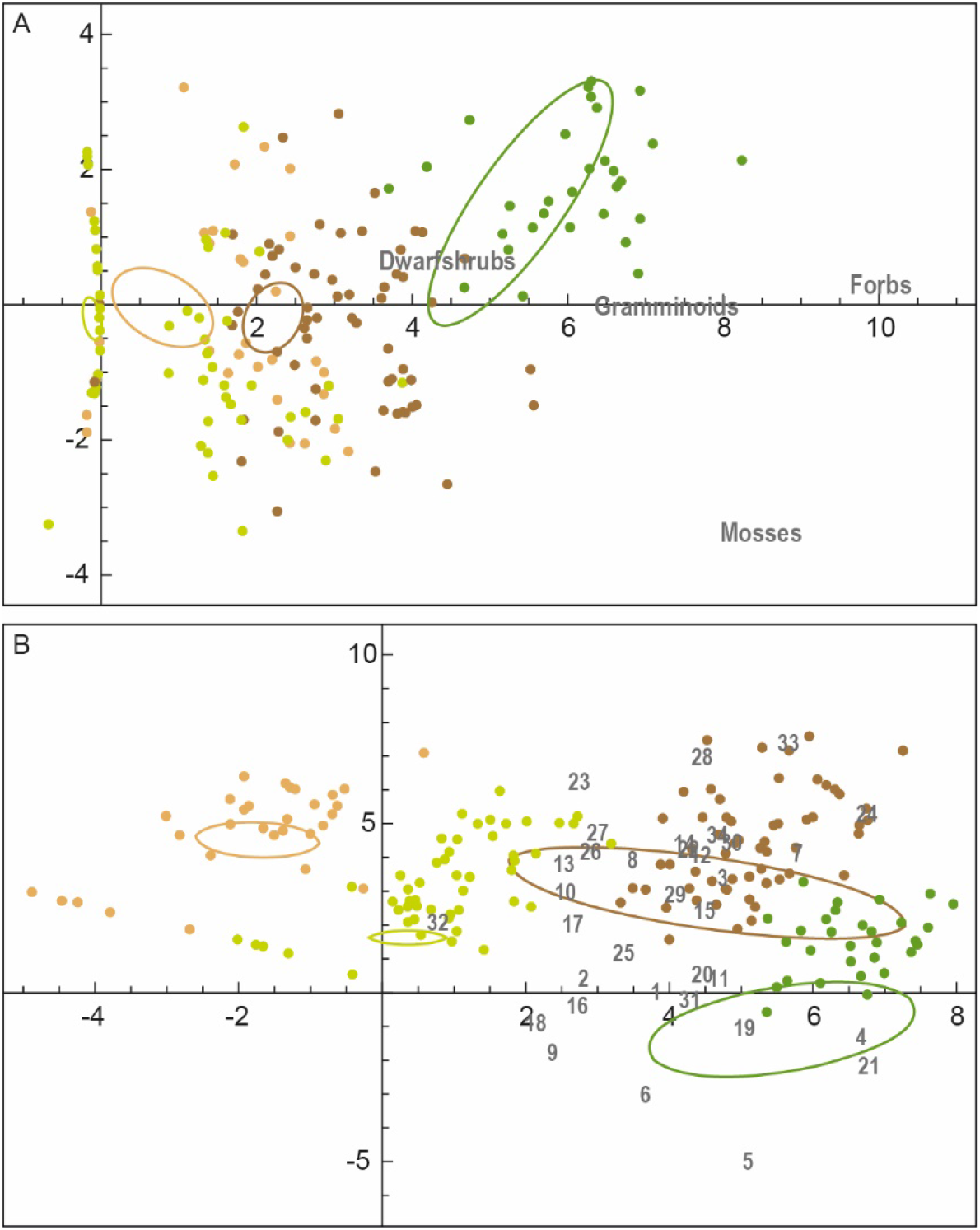
Map of latent variables, posterior means of (***α**_i_* + ***γ**_i_* + ***z**_i_*) and 95% credibility areas of the effect of fixed factors (management), brown: burned, light green: harvested, dark brown: unmanaged, dark green: grazed. Mean species coefficients, ***θ**_i_* for each cover class (species) are shown. A: four aggregated species classes. B: All species, 1: *Eriophorum angustifolium*, 2: *Calamagrostis epigejos*, 3: *Trichophorum cespitosum*, 4: *Danthonia decumbens*, 5: *Agrostis capillaris*, 6: *Nardus stricta*, 7: *Festuca ovina*, 8: *Deschampsia flexuosa*, 9: *Festuca rubra*, 10: *Molinium caerulea*, 11: *Poa compressa*, 12: *Carex pilulifera*, 13: *Carex arenaria*, 14: *Carex panacea*, 15: *Carex nigra*, 16: *Luzula campestris*, 17: *Galium saxatile*, 18: *Hieracium pilosella*, 19: *Potentilla erecta*, 20: *Rumex acetosella*, 21: *Hypochoeris radicata*, 22: *Vaccinium vitis-idaea*, 23: *Calluna vulgaris*, 24: *Empetrum nigrum*, 25: *Genista anglica*, 26: *Erica tetralix*, 27: *Dicranum scoparium*, 28: *Pseudoscleropodium purum*, 29: *Pleurozium schreberi*, 30: *Bryum* subsp., 31: *Sphagnum compactum*, 32: *Campylopus introflexus*, 33: *Hypnum cupressiforme*, 34: *Cladonia rangiferina*.

The main results are shown in Fig. 2, where the mean of the posterior distribution of (***α**_i_* + ***γ**_i_* + ***z**_i_*) vectors are displayed for each plot for the aggregated species classes and all species, respectively. The distance between plots indicates the difference in the cover of the species, where the scale is relative to the unit standard deviation of the residual latent variables, i.e. the variation in species cover among plots that could not be explained by the fixed or random factors.

The 95% credibility areas of the effect of fixed factors, i.e. the different management regimes, are shown by colored ovals in Fig. 2. Since no ovals overlap, we concluded that the four different management regimes lead to characteristic heathland plant communities, which on average were distinct from each other. In particular, grazing has a large effect on the composition of the heathland vegetation, whereas the effects of the other three management regimes were comparably more similar (Fig. 2). This is further demonstrated in Fig. 1, where grazing leads to a shift in vegetation cover from a high dwarf-shrub dominance towards forb and graminoid covered heathland. Inclusion of all species in the model still yielded a distinct plant species composition with non-overlapping credibility intervals. However, the relative positions of burned and harvested plots were shifted, indicating that inference of management affects species composition depending on the taxonomic resolution (functional groups vs. all species). Furthermore, increasing the taxonomic resolution leads to higher within treatment variation in the unmanaged plots compared to the aggregated dataset (Fig 2b, larger credibility area).

The scale of the random effects, i.e. the effect of site, is shown in Table 1. Given that the scale of the random effect is relative to the unit standard deviation of the residual latent variables, then we conclude that the displacement due to sites tended to be significantly larger than the variation among the residual latent variables (Table 1, *σ_2_* is significantly larger than one for both cases). This indicated a relatively large influence of site-specific species pools. There was also a significant positive correlation between the two dimensions of the random effects when all species were analyzed (Table 1).

In both cases, the estimated posterior distribution of *δ* was dominated by the chosen lower limit of the prior distribution of the parameter (0.01), and the degree of spatial aggregation could not be distinguished from zero in a statistical sense, i.e. random expectations.

## Discussion

Over the past five years, there has been an increasing interest in model-based approaches for the statistical modelling of the joint distribution of multi-species abundances (e.g. Clark et al. 2014, Warton et al. 2015, Warton et al. 2016, Ovaskainen et al. 2017). The current study is an example reflecting this important trend in community ecology, as the importance of how species abundance data are distributed, sampled, and driven by a complex interplay of different ecological processes is given increasing recognition. Plant species are sedentary, and in many terrestrial ecosystems plants dominate the ground cover, leaving few bare patches. These characteristic features of plant growth have important consequences for the joint distribution of plant species cover, such as large spatial aggregation among single species and relatively strong negative correlation among them. To model these features, the Dirichlet-multinomial distribution has previously been suggested as a suitable candidate distribution for the joint species distribution of pin-point plant cover data (Damgaard 2015, Damgaard et al. 2017, Damgaard 2019). The Dirichlet-multinomial distribution has markedly different distributional properties than both the Bernoulli distribution and the negative binomial distribution, which till now have typically dominated the development of model-based ordination methods and software (e.g. Niku et al. 2019).

In the suggested reparametrized Dirichlet-multinomial distribution, the spatial aggregation among plant species is modelled by the parameter *δ*, which increases with the within-site spatial variation in cover, i.e. the degree of spatial aggregation. In the present case, the degree of spatial aggregation could not be distinguished from random expectations, and it was concluded that the observed covariation pattern among the species was adequately modelled by the residual latent variables. Consequently, we suggest to test whether the more simple multinomial distribution is suitable in future model-based ordination of pin-point cover data.

Unlike in Hui et al. (2015) and Warton et al. (2015), among others, both the fixed and random factors are here modelled as p-dimensional vectors and added to the latent variables before the inner product with the species-specific coefficients. This means that the estimated displacement effects of management and sites are estimated and statistical inference performed in the same *p*-dimensional latent ordination space. When *p* = 2 in particular, the estimated average displacements of e.g. the different management regimes may be shown in the plane as ovals of 95% credibility areas. Such plots, we find, are an illustrative way to show the effects of different treatments on the species composition. This is in contrast to most latent variable models, which consider the sample scores as random effects and make prior distributional assumptions on them without incorporating both random and fixed effects in the same model framework. Naturally, the suggested parameterization with fixed and random effects may also be applicable for count data or absence-presence data.

Not surprisingly, in our case study the interpretation of the fixed effects is sensitive to taxonomic resolution of the dataset. This was evident when comparing Fig 2a and b. If data was aggregated into functional groups (Fig. 2a), it could be concluded that management abandonment leads to relatively homogenous plant communities. However, the model based on species level resolution (Fig. 2b) showed that the variation within the unmanaged plots is high. Regardless of taxonomic resolution, both models exhibited clear evidence that a varied management regime with different management methods produces an overall heterogeneous plant species composition. Hence, interpretation of fixed effects in this model-based framework may provide important information for heathland conservation at both the level of plant functional groups and at the species level resolution.

If *p* >2, then the 95% credibility regions, which illustrate the mean effect of the fixed and random factors, must be imagined as a *p*-dimensional ellipsoidal shape. We expect that the statistical power to separate the effects of the different fixed factors on species composition will increase with the dimension of the latent variable used in the ordination. However, this preliminary notion needs to be explored in a more systematic way. Finally, the chosen estimation method is relatively slow, and in cases with many plots and many species faster estimation procedures may be needed and should be developed in the future.

## Acknowledgements

We thank K. E. Nielsen for field help providing pin-point data. CFD and RRH both acknowledge the funding provided by Aage V. Jensen foundation. FKCH acknowledges the funding provided by two grants from the Australian Research Council

## Electronic supplements

## Appendix A: Details of the case study

A map of Jutland showing the location and size of the heathlands investigated in the study. Inset figure shows the location of the selected plot within the management parcel. Inset figure shows the experimental design within each plot. The squares show an example of the randomised locations for pin-point analyses.

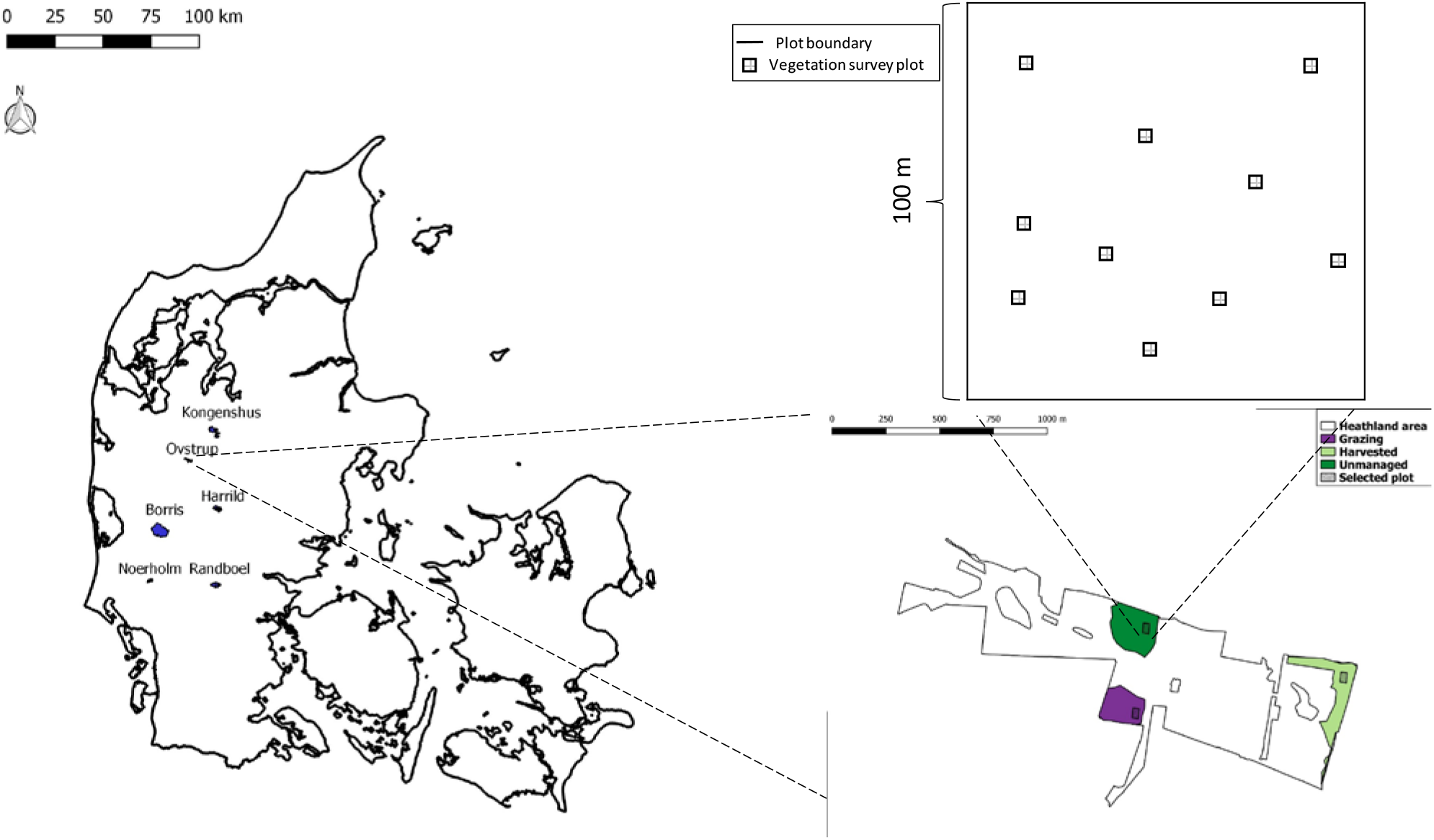

An overview of the selected sites and the management carried out in the selected plots. The number under burn, harvest, and unmanaged is the number of years since the management was last applied. For grazing, the number is the number of years it has been grazed. Frequency denotes how often the management is performed.

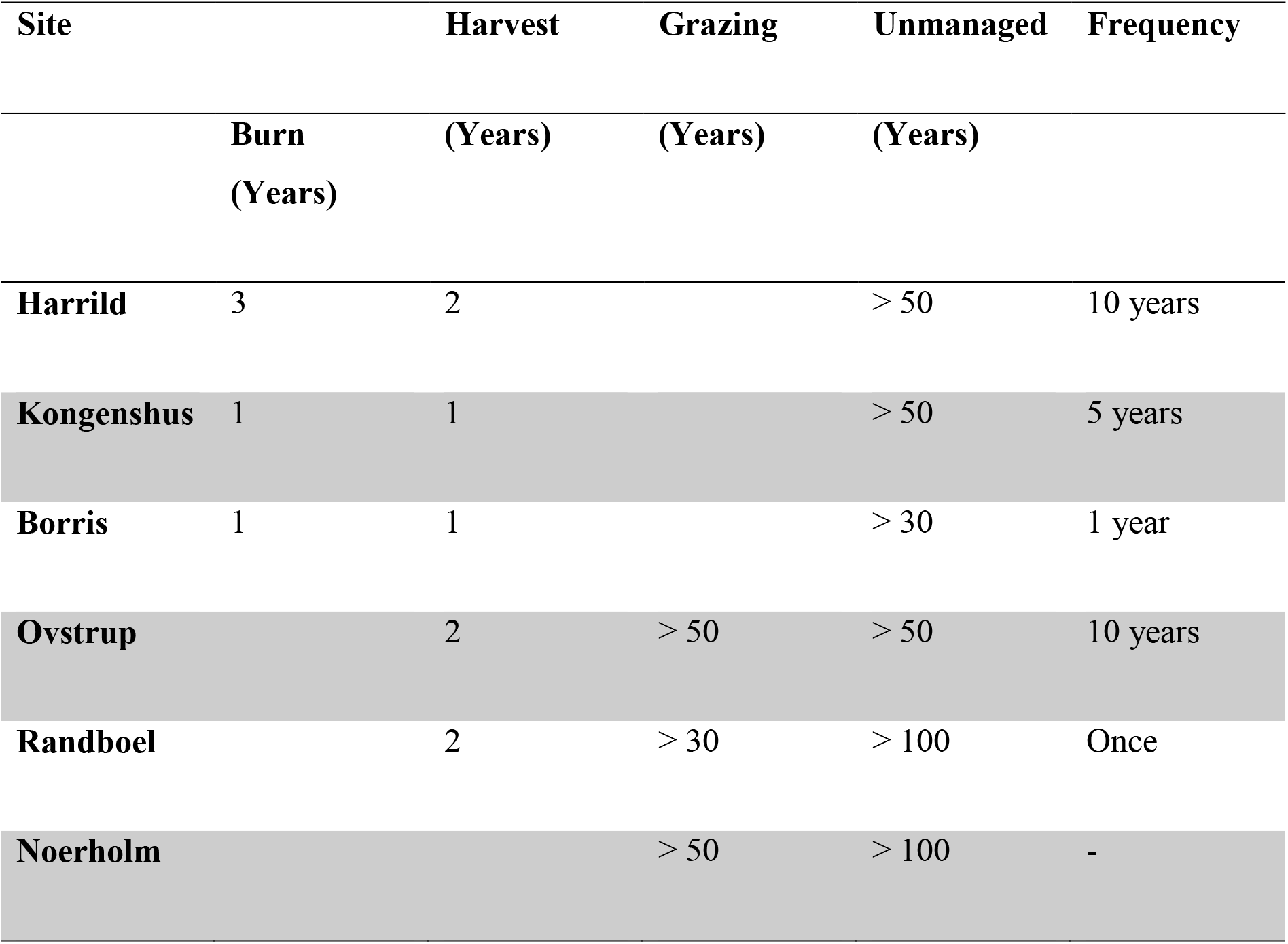

## Appendix B: Mathematica notebook

